# Structural basis for mTORC1-dependent regulation of the lysosomal and autophagic transcription factor TFEB

**DOI:** 10.1101/2022.09.12.507619

**Authors:** Zhicheng Cui, Gennaro Napolitano, Mariana E. G. de Araujo, Alessandra Esposito, Jlenia Monfregola, Lukas A. Huber, Andrea Ballabio, James H. Hurley

**Affiliations:** Department of Molecular and Cell Biology, University of California Berkeley; Berkeley CA 94720, USA; California Institute for Quantitative Biosciences, University of California, Berkeley, CA, 94720, USA; Telethon Institute of Genetics and Medicine (TIGEM), Naples, Italy; Medical Genetics Unit, Department of Medical and Translational Science, Federico II University, Naples, Italy; Institute of Cell Biology, Biocenter, Medical University of Innsbruck, Innsbruck, Austria; Department of Molecular and Human Genetics, Baylor College of Medicine, Houston, TX, USA; Jan and Dan Duncan Neurological Research Institute, Texas Children’s Hospital, Houston, TX, USA; SSM School for Advanced Studies, Federico II University, Naples, Italy; Helen Wills Neuroscience Institute, University of California, Berkeley, Berkeley, CA 94720, USA

## Abstract

The transcription factor TFEB is a master regulator of lysosomal biogenesis and autophagy. The phosphorylation of TFEB by the mechanistic target of rapamycin complex 1 (mTORC1) is unique in its mTORC1 substrate recruitment mechanism, which is strictly dependent on the amino-acid-mediated activation of the RagC GAP FLCN. TFEB lacks the TOR signaling (TOS) motif responsible for the recruitment of other mTORC1 substrates. We used cryo-electron microscopy (cryo-EM) to determine the structure of TFEB as presented to mTORC1 for phosphorylation. Two full Rag-Ragulator complexes present each molecule of TFEB to the mTOR active site. One Rag-Ragulator complex is bound to Raptor in the canonical mode seen previously in the absence of TFEB. A second Rag-Ragulator complex (non-canonical) docks onto the first via a RagC GDP-dependent contact with the second Ragulator complex. The non-canonical Rag dimer binds the first helix of TFEB in a RagC^GDP^-dependent aspartate clamp in the cleft between the Rag G domains. Mutation of the clamp drives TFEB constitutively into the nucleus whilst having no effect on mTORC1 localization. The remainder of the 108-amino acid TFEB docking domain winds around Raptor and then back to RagA. This structure presents the phosphorylatable Ser residues of TFEB to the mTORC1 active site in a suitable geometry for their phosphorylation. The double use of RagC GDP contacts in both Rag dimers explains the strong dependence of TFEB phosphorylation on FLCN and the RagC GDP state.

## Introduction

The transcription of genes responsible for lysosome biogenesis and autophagy, known as the coordinated lysosomal expression and regulation (CLEAR) network, is under the control of transcription factor EB (TFEB) ^1^. TFEB is one of four members of the microphthalmia (MiT) family of basic helix-loop-helix leucine zipper (bHLH-Zip) transcription factors ^2^. Overexpression of TFEB promotes degradation of long-lived proteins ^3^, lipid droplets ^4^, and damaged mitochondria ^5^, and can induce lysosomal exocytosis ^6^. Indeed, data from cellular and mouse models show that TFEB activation increases autophagic and lysosomal clearance capacity, and is therefore a potential therapeutic target for the treatment of lysosomal storage disorders^7,8^ and neurodegenerative diseases involving damaged organelles and accumulation of protein aggregates ^9–12^. The latter include Parkinson’s Disease and Alzheimer’s Disease. TFEB is regulated by cellular nutrient status through its multisite phosphorylation at several serine residues, including Ser 122, Ser142 and Ser211, by the mechanistic target of rapamycin complex 1 (mTORC1) under nutrient-replete conditions ^13–16^. Hierarchical phosphorylation of these sites allows TFEB cytosolic retention and inactivation ^14,15,17^. mTORC1 is recruited to the lysosomal membrane for activation by the Rag GTPases, which are heterodimers composed of Rag A or B bound to Rag C or D ^18–20^. The pentameric Ragulator/Lamtor complex, composed of Lamtor1-5 proteins, is a scaffold that anchors the Rags to the lysosomal membrane via myristoyl and palmitoyl posttranslational modifications of its Lamtor1 subunit ^21^.

The cryo-EM structures of Rag dimers bound to mTORC1^22^ or its Raptor subunit^23^ showed that RagA^GTP^ extensively contacts Raptor. Despite the importance of RagC^GDP^ in mTORC1 physiology, these structures also showed RagC^GDP^ interacts with Raptor to a lesser degree and without stringent dependence on the RagC nucleotide state. The tumor suppressor FLCN is the GTPase activating protein (GAP) for RagC ^24^. FLCN activity is required for the phosphorylation of TFEB, but not other mTORC1 substrates ^25^. FLCN is maintained in the inactive lysosomal FLCN complex (LFC) in amino acid starvation ^26^. FLCN is reactivated under amino acid replete condition when the LFC is destabilized by the amino acid transporter SLC38A9 ^27^. TFEB phosphoregulation accounts for the tumor suppressor function of FLCN in Birt-Hogg-Dubé (BHD) syndrome ^25^. TFEB lacks the TOR signaling (TOS) motif found in other mTORC1 substrates ^28^, which enables presentation of substrates to the catalytic subunit by Raptor ^29^. Instead, TFEB was shown to interact with the Rag GTPases^30^, which serve as a substrate recruitment mechanism that allows TFEB phosphorylation by mTORC1 ^25^. We therefore hypothesized that a unique structural platform directly involving RagC^GDP^ might be responsible for selectively presenting TFEB as a substrate of mTORC1. We set out to test the hypothesis by reconstituting and determining the structure of the complex.

### Reconstitution and cryo-EM structure of the Raptor-TFEB-Rag-Ragulator complex

A complex of TFEB-RagA-RagC was obtained by co-expression of full-length TFEB (R245-247A, S211A) and Rag GTPases (RagA^Q66L^, RagC^S75N^) in human embryonic kidney (HEK)-293F GnTI-cells, and found it to be stable under size exclusion chromatography (Extended Data Fig. 1a,b). Mutations in the nuclear localization signal (NLS) of TFEB (R245A, R246A, R247A) were introduced to prevent nuclear translocation^30^ during expression in HEK cells. We also introduced the S211A mutation in TFEB because it had been reported to stabilize TFEB association with the Rags in cells ^30^. The mutations RagA (Q66L) and RagC (S75N) were incorporated to promote the active configuration of Rag GTPases (RagA^GTP^:RagC^GDP^). We reconstituted the TFEB-Rag complex with purified Ragulator complex and solved the cryo-EM structure (Extended Data Fig. 1a, c-e). However, no TFEB density was observed (Extended Data Fig. 1f), indicating that additional interactions were required to generate a complex stable enough for vitrification. Previous structures ^22,23^ suggested that Raptor would be structurally proximal to the TFEB binding site. We repeated the reconstitution in the presence of purified Raptor, assembled a Raptor-TFEB-Rag-Ragulator complex (Fig. 1a), and determined its cryo-EM structure to an overall 3.1-Å resolution (Extended Data Fig. 2a, d, Table 1).

**Fig. 1.**
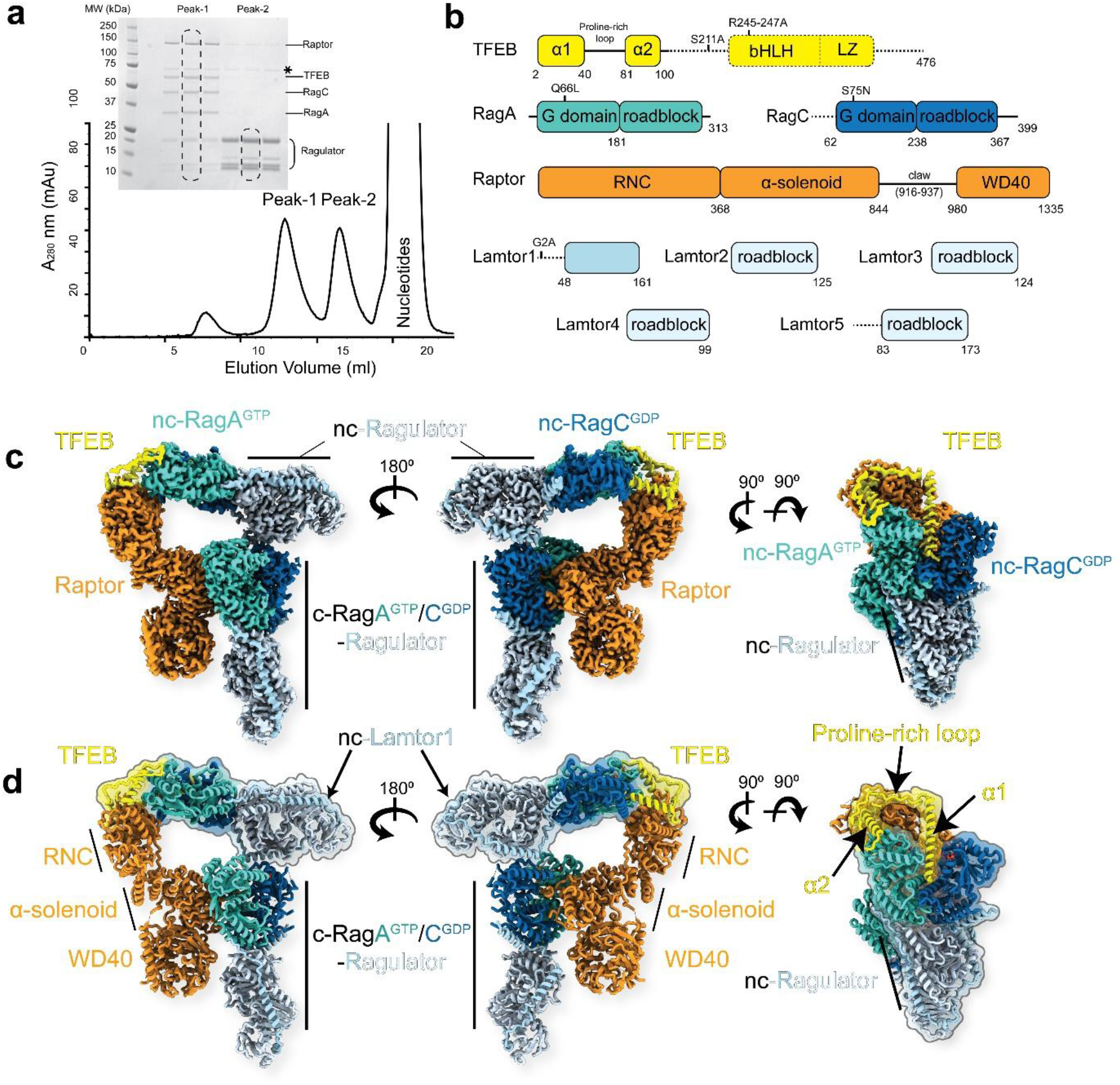
Reconstitution and structural determination of the Raptor-TFEB-Rag-Ragulator complex. **a**, Size exclusion chromatography and SDS-polyacrylamide gel electrophoresis of assembled Raptor-TFEB-Rag-Ragulator complex. Peak-1 corresponds to the fully assembled complex, and peak-2 represents Ragulator alone. All the corresponding bands are labeled, the asterisk indicates HSP70 contamination. **b**, Domain arrangement of all the subunits in the complex. Unresolved domains are indicated by dash lines. **c**, A composite cryo-EM density map of the complex, assembled from three focused-refinement maps (Raptor, c-RagA^GTP^/RagC^GDP^-Ragulator, and TFEB-nc-RagA^GTP^/RagC^GDP^-Ragulator). Different contour levels were used for optimal visualization using UCSF ChimeraX ^51^. **d**, Atomic model of the complex in the same viewing orientation as in (**c**). Domains for Raptor and TFEB are labeled. The TFEB-nc-RagA^GTP^/RagC^GDP^-Ragulator is highlighted in an outline.

The structure of Raptor-TFEB-Rag-Ragulator revealed a 2:1 stoichiometry for Rag-Ragulator with respect to Raptor, as compared to 1:1 observed in the structures of mTORC1/Raptor-Rag-Ragulator ^22,23^ (Fig. 1c, d). We refer to the Rag-Ragulator module that binds to Raptor as previously reported as the “canonical” (c) Rag-Ragulator, and the second module as the “non-canonical” (nc) Rag-Ragulator. Further local refinement of Raptor, canonical Rag-Ragulator, and TFEB-nc-Rag-Ragulator yielded cryo-EM maps to the resolutions of 2.8 Å, 2.9 Å, and 2.9 Å, respectively, which allowed us to build accurate atomic models (Extended Data Fig. 2a, d). The cryo-EM density of TFEB was clearly visualized and corresponds to the residues 2-105 (Extended Data Fig. 3a). The N- and C-termini of this segment are helical and referred to as α1 and α2, and they are connected to a Pro-rich loop. The TFEB-nc-Rag-Ragulator spans the Raptor and the canonical Rag dimer, forming a closed triangle structure with extensive contacts at both ends (Fig. 1d, Extended Data Fig. 4). Both Rag heterodimers are in active states, based on the high-resolution cryo-EM density of the corresponding nucleotide (Extended Data Fig. 3b).

### Interactions between TFEB and active Rag GTPases

The structure showed that TFEB residues 2-105 were ordered, suggesting that these residues were both necessary and sufficient to form a stable complex with active Rag GTPases, and we confirmed that a slightly longer TFEB 1-109 construct was competent to form such a complex *in vitro* (Extended Data Fig. 5). The first 40 residues of TFEB form a long helix (α1) and occupy the cleft between the two G domains of the TFEB-specific dimer of RagA^GTP^ and RagC^GDP^ (Fig. 2a). Residues 2-18 of TFEB α1 are embedded in the inter-Rag G domain cleft and form a ~ 570 Å interaction interface with the RagC^GDP^ G domain (Fig. 2b). The N-terminus of TFEB α1 sits directly on top of the α8 of RagC^GDP^ roadblock domain, which is, in turn, at the dimerization interface of Rag heterodimer. TFEB helix α1 is thus clamped within the G domain cleft by hydrogen bonds between Asp^294^ of RagC^GDP^ and the backbone of Arg^4^ and Ile^5^ of TFEB and salt bridges between Asp^290^ of RagC^GDP^ and Arg^4^ and Arg^8^ of TFEB (Fig. 2g). The α1 of TFEB also contacts RagC^GDP^ via hydrophobic interactions between Leu^7^, Leu^11^ of TFEB and Val^80^, Ile^220^ of RagC^GDP^, and between the side chain of Gln^15^ of TFEB and Tyr^221^ of RagC^GDP^ (Fig. 2b). A stacking interaction is also present between the carbon chain of Arg^13^ of TFEB and Trp^165^ of RagA^GTP^. There are fewer specific contacts between TFEB α1 and RagA^GTP^, primarily through hydrophobic interactions between Ile^5^, Met^9^ of TFEB and Ile^234^ of RagA^GTP^. These data show how the TFEB N-terminus is clamped between the Rag G domains in a strictly RagC^GDP^-dependent manner.

**Fig. 2.**
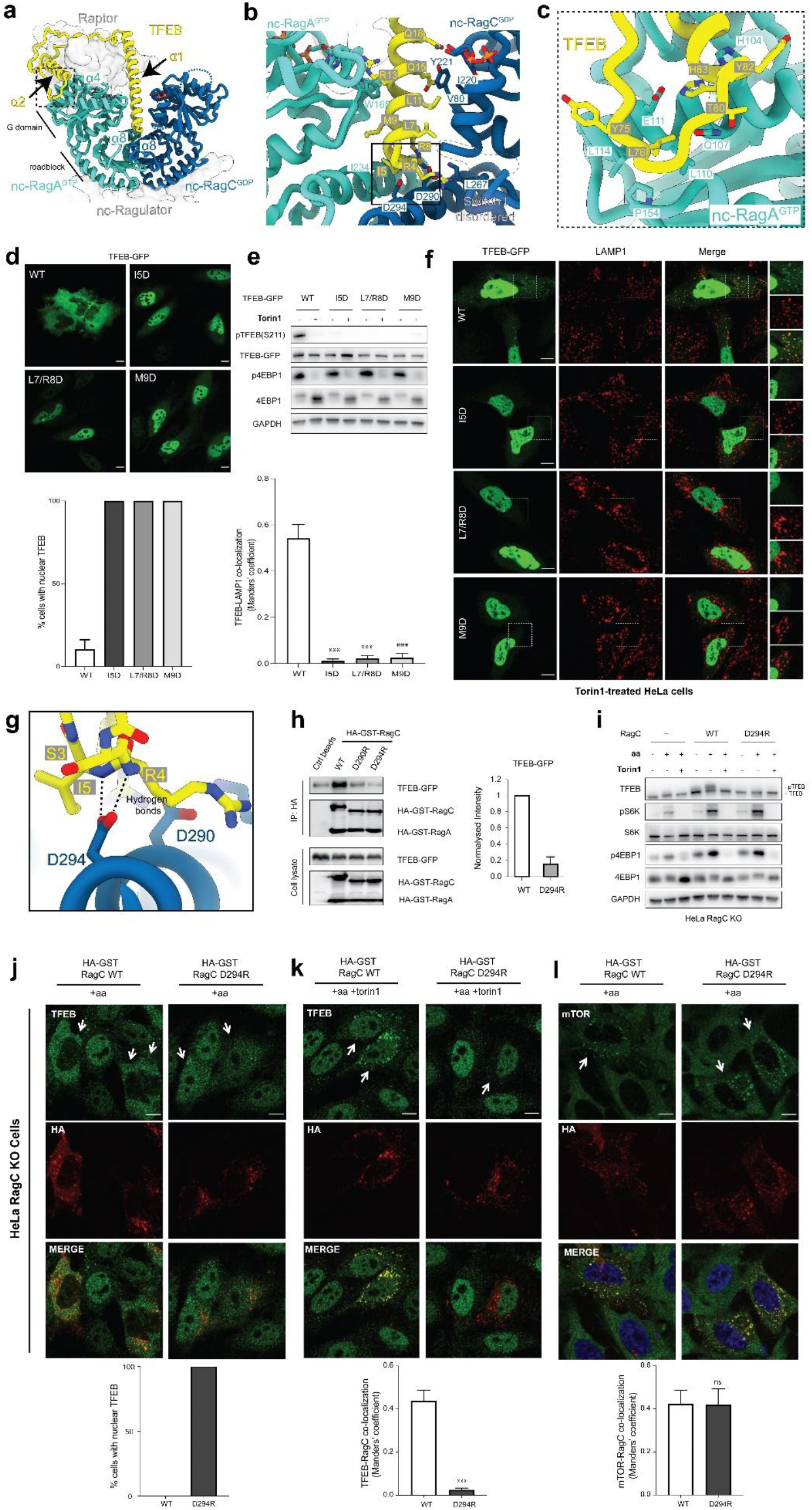
TFEB recruitment by active Rag GTPases. **a**, Overall interaction between TFEB and nc-Rag GTPases is shown as in ribbon models from the front view. Raptor and nc-Ragulator are shown as transparent surfaces. Disordered switches I and II of nc-RagC^GDP^ are shown in dash lines. **b**, Interactions between TFEB and inter-Rag G domains at the dimer interface. **c**, Close-up view of the interaction between TFEB and outer-G domain of RagA^GTP^ as outlined in (**a**). **d**, Cells expressing wild type or mutant TFEB-GFP were analyzed by immunofluorescence and quantified to calculate the percentage of cells showing TFEB nuclear localization. Scale bar, 10 μm. **e**, Representative immunoblotting of HeLa cells expressing wild type or mutant TFEB-GFP. **f**, Confocal microscopy analysis of Torin1-treated HeLa cells described in (**e**), showing TFEB and lysosome co-localization. Scale bar, 10 μm. Quantifications of TFEB-LAMP1 colocalization are shown on the left. **g**, Close-up view of the interaction between TFEB N-terminus and α8 of RagC^GDP^ as outlined in (**b**). Hydrogen bonds are labeled and indicated with dash lines. **h**, HEK293A cells stably expressing TFEB-GFP were transfected with the indicated constructs. Cells were then lysed, incubated with HA beads and analyzed by immunoblotting (replicated three times). HA, haemagglutinin; GST, glutathione *S*-transferase; IP, immunoprecipitation. **i**, Representative immunoblotting of RagC-knockout HeLa cells transfected with control vector (empty) or wild type RagC or RagC(D294R). Cells were subjected to amino acid starvation and refeeding, in the presence or absence of 250 nM torin. **j-l**, Cells as in (**i**) were analyzed by immunofluorescence and quantified to calculate the percentage of the cells showing TFEB nuclear localization (**j**), TFEB-RagC colocalization (**k**), and mTOR-TFEB colocalization (**l**). Scale bar, 10 μm. TFEB and mTOR were stained with corresponding antibodies. Colocalization was calculated with Manders’ colocalization coefficient.

To validate the functional role of the TFEB N-terminal clamp, both the N-terminal residues of TFEB and Asp^290^ or Asp^294^ of RagC were mutated. TFEB mutants I5D, L7D/R8D, and M9D were reconstituted into HeLa cells as GFP fusions, and expressed at essentially equal levels (Fig. 2d, e). Cytosolic localization and Ser211 phosphorylation of wild-type TFEB was observed robustly in the absence of torin, as expected. Instead, all three of the mutants showed constitutively nuclear localization and defective Ser211 phosphorylation, even in the absence of torin (Fig. 2d). Lysosomal localization, which is observed for wild-type TFEB in torin-treated cells, was essentially abolished for all of the mutants (Fig. 2f), confirming the functional requirement for these extreme N-terminal residues of TFEB in the G domain clamp. Next we used a previously established HEK293A cell line stably expressing TFEB-GFP to validate the biochemical analysis. The interaction between TFEB-GFP and the transiently transfected Rags was significantly impaired in both RagC^D290R^ and RagC^D294R^ in comparison to wild-type RagC (Fig. 2h). Moreover, expression of RagC^D294R^ in RagC knockout cells prevented TFEB phosphorylation in amino acid replete cells, even in the absence of torin (Fig. 2i). However, the phosphorylation of S6K and 4E-BP1 was supported normally, indicating that these TOS motif substrates do not require the RagC clamp. RagC^D294R^ expression supported mTORC1 localization to lysosomes in amino acid replete conditions (Fig. 2l), but not cytosolic and lysosomal localization of TFEB in the absence and presence of torin, respectively (Fig. 2j, k). These data support that the RagC G domain clamp uniquely regulates TFEB phosphorylation.

TFEB only interacts with the active RagC^GDP^-containing Rag dimer. Alignment of the TFEB bound active Rag GTPases and inactive Rag GTPases (RagA^GDP^:RagC^GTP^) ^26,27,31^ based on the α8 of RagC roadblock domains indicated that the switch I of RagC^GTP^ sterically clashes with the α1 of TFEB, therefore precluding TFEB binding (Extended Data Fig. 6). Notably, a previously unknown interaction interface between residues 76-83 of TFEB and α4 of RagA^GTP^ is also revealed in our structure (Fig 2c). Leu^76^ of TFEB inserts into a hydrophobic pocket formed by Leu^110^, Leu^114^, Pro^154^, and Leu^155^ of RagA^GTP^. A salt bridge between His^84^ of TFEB and Glu^111^ of RagA^GTP^ is also observed. Alignment of the TFEB bound active RagA^GTP^ and inactive RagA^GDP^ based on the α4 of RagA G domain revealed apparently insignificant structural differences in the TFEB binding region (Extended Data Fig. 7), suggesting the nucleotide loading state of RagA does not impose selectivity toward TFEB-binding at the direct RagA-TFEB interface. However, the wide opening conformation of G domains in the inactive Rag GTPases (RagA^GDP^:RagC^GTP^) and the fact that GTP loaded RagC sterically hinders the binding of TFEB N terminus to the G domain clamp, effectively prevent the interaction between TFEB and the Rag heterodimer in the inactive state (Extended Data Fig. 6). Thus, while RagC^GDP^ is uniquely important for TFEB but not other substrates, RagA^GTP^ is important for TFEB phosphorylation just as for all known mTORC1 substrates.

### Bridging two Rag-Ragulator complexes

The TFEB-nc-Rag-Ragulator complex is stabilized by interactions at both ends (Fig. 3a, 4a). On one end, the Pro-loop and α2 of TFEB bridge the Raptor N-terminal conserved (RNC) domain and the ordered switch I of nc-RagA^GTP^ (Fig. 3b-d). Since the switch I of RagA is disordered in the GDP-bound state ^26,27^, it emphasizes the importance of nc-Rag GTPases being in the active state (RagA^GTP^:RagC^GDP^). The residues Thr^50^, Pro^51^, Ala^52^, and Ile^53^ of TFEB cover a hydrophobic patch on the RNC domain, formed by residues Pro^73^, Pro^156^, Trp^165^, Try^174^, Ile^175^, and Pro^176^ (Fig. 3e). The contact between Raptor and nc-RagA^GTP^ is maintained by hydrophobic interaction between Leu^133^, Leu^137^ of Raptor and Ile^43^ of nc-RagA^GTP^ (Fig. 3d).

**Fig. 3.**
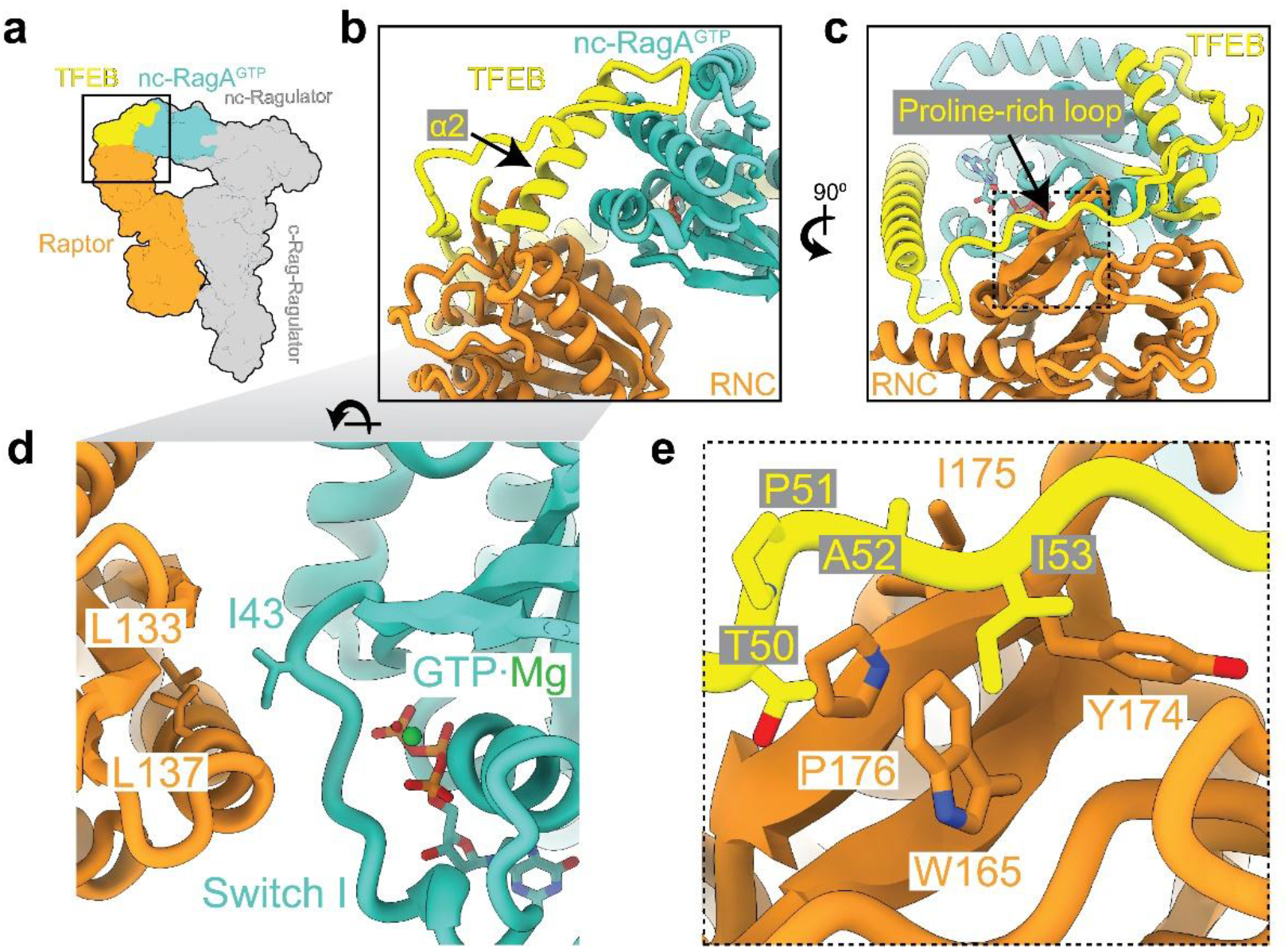
TFEB bridges non-canonical Rag-Ragulator through its interaction with Raptor. **a**, Cartoon representation that highlights the interacting subunits at the end with TFEB. **b**, Ribbon model showing the interactions among TFEB, Raptor and nc-RagA^GTP^. TFEB bridges the interaction between Raptor and nc-RagA^GTP^ through its Pro-rich loop and α2 region. **c**, 90°-rotated view of (**b**) shows the interaction between the Pro-rich loop of TFEB and RNC domain. **d**, Closeup view of the interaction between RNC domain and nc-RagA^GTP^. Ordered switch I of nc-RagA^GTP^ facilitates its interaction with the RNC domain. **e**, Close-up view as outlined in (**c**) shows the residues responsible for the hydrophobic interaction between Pro-rich loop of TFEB and RNC domain.

**Fig. 4.**
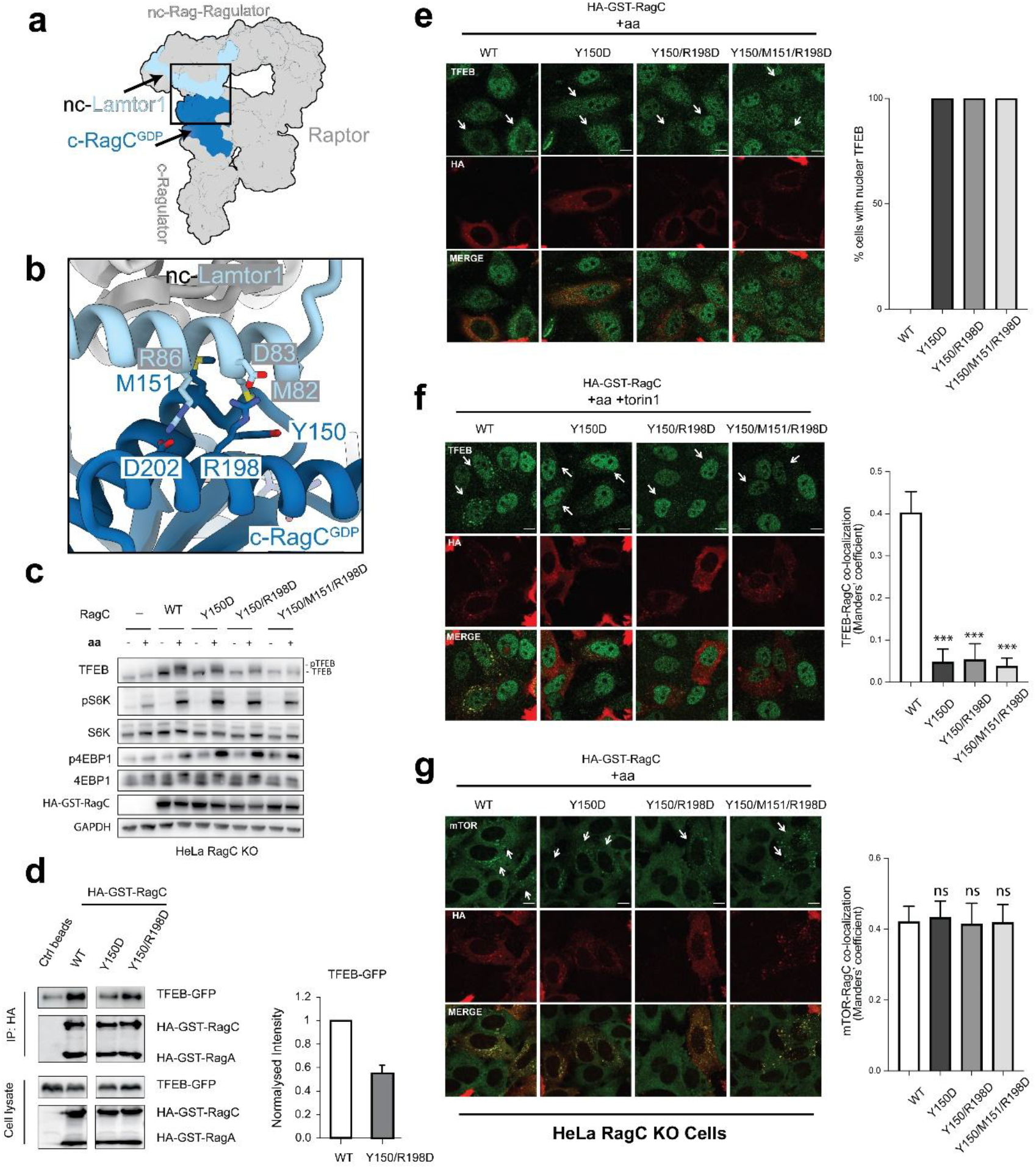
The non-canonical Lamtor1 and canonical RagC^GDP^ contact is required for TFEB phosphorylation. **a**, Cartoon representation that highlights the interacting subunits at end with nc-Ragulator. **d**, Close-up view as outlined in (**a**) shows the residues responsible interaction between nc-Lamtor1 and c-RagC^GDP^. **c**, Representative immunoblotting of RagC-knockout HeLa cells transfected with control vector (empty) or wild type RagC or RagC mutants (Y150, Y150/R198D, or Y150/M151/R198D). Cells were then subjected to amino acid starvation and refeeding, in the presence or absence of 250 nM torin. **d**, Cell lysates from HEK293A cells stably expressing TFEB-GFP were incubated with HA beads and analyzed by immunoblotting. **e-g**, Cells as in (**c**) were analyzed by immunofluorescence and quantified to calculate the percentage of the cells showing TFEB nuclear localization (**e**), TFEB-RagC colocalization (**f**), and mTOR-TFEB colocalization (**g**). Scale bar, 10 μm. TFEB and mTOR were stained with corresponding antibodies. Colocalization was calculated with Manders’ colocalization coefficient.

On the other side of the interface, the Lamtor1 subunit of the non-canonical Ragulator makes a close contact with the G domain of canonical RagC^GDP^ (Fig. 4a). The α2 of Lamtor1 resides on top of α4 and α5 helices of RagC, stabilized by Met^82^ of Lamtor1 inserting into the hydrophobic pocket formed by Tyr^150^, Met^151^, Leu^154^ of RagC^GDP^ (Fig. 4b). Salt bridges are formed between Asp^83^, Arg^86^ of Lamtor1 and Arg^198^, Asp^202^ of RagC^GDP^, respectively. These data suggest that the RagC^GDP^ state in the canonical Rag dimer is important for nc-Lamtor1 interaction and stabilization of the nc-Rag-Ragulator binding, thus heightening the sensitivity of the entire assembly to the RagC nucleotide state and thus the dependency of TFEB phosphorylation on FLCN.

To validate the role of the c-RagC and nc-Ragulator interaction, RagC residues Tyr^150^, Met^151^, and Arg^198^ were mutated and reconstituted in RagC-deleted HeLa cells. The constructs RagC^Y150D^, RagC^Y150D/R198D^, and RagC^Y150D/M151D/R198D^ all supported normal S6K and 4E-BP1 phosphorylation in amino acid replete conditions, but showed reduced or no TFEB Ser211 phosphorylation under these same conditions (Fig. 4c). Consistently, only wild-type RagC restored TFEB cytosolic localization in RagC-KO cells (Fig. 4e). Furthermore, in contrast to wild-type RagC, none of the RagC mutants was able to restore TFEB lysosomal localization in torin-treated RagC-KO cells, consistent with the impaired ability of TFEB to form a stable lysosomal complex with mutant RagC and Ragulator (Fig. 4f). Consistent with these functional data, we observed a reduced interaction between TFEB and the transiently transfected RagC^Y150D^ and RagC^Y150D/R198D^ (Fig. 4d). Taken together, these data show that not only is the G domain clamp in the nc-Rag dimer essential for TFEB recruitment to lysosomes, but the c-Rag to nc-Ragulator contact as well. Wildtype and all three RagC mutants were similarly able to restore mTOR lysosomal localization in RagC-KO cells (Fig. 4g), consistent with a selective role for this interaction in regulation the phosphorylation of TFEB but not other mTORC1 substrates.

### Cryo-EM structure of the mTORC1-TFEB-Rag-Ragulator megacomplex

To understand how TFEB is phosphorylated by mTORC1 as presented by the Raptor-Rag-Ragulator complex, we reconstituted mTORC1-TFEB-Rag-Ragulator megacomplex and determined its structure by cryo-EM. Two major populations of the megacomplex were resolved, showing that either one or two copies of TFEB are present on mTORC1 (Extended Data Fig. 8a). The reconstruction in C2 symmetry for the mTORC1 with two copies of TFEB resulted in a 3.7-Å resolution cryo-EM map. Further symmetry expansion and local refinement of the asymmetric unit improved the resolution to 3.2 Å (Extended Data Fig. 8e, Table 1). A composite map was generated by superimposing the local reconstructions to the C2 symmetric reconstruction (Fig. 5a). We then built the atomic model for the entire complex, which contains mTOR, Raptor, mLST8, TFEB, active Rag GTPases, Ragulator with a stoichiometry of 2:2:2:2:4:4, containing a total of 36 polypeptide chains (Fig. 5b, Movie 1).

**Fig. 5.**
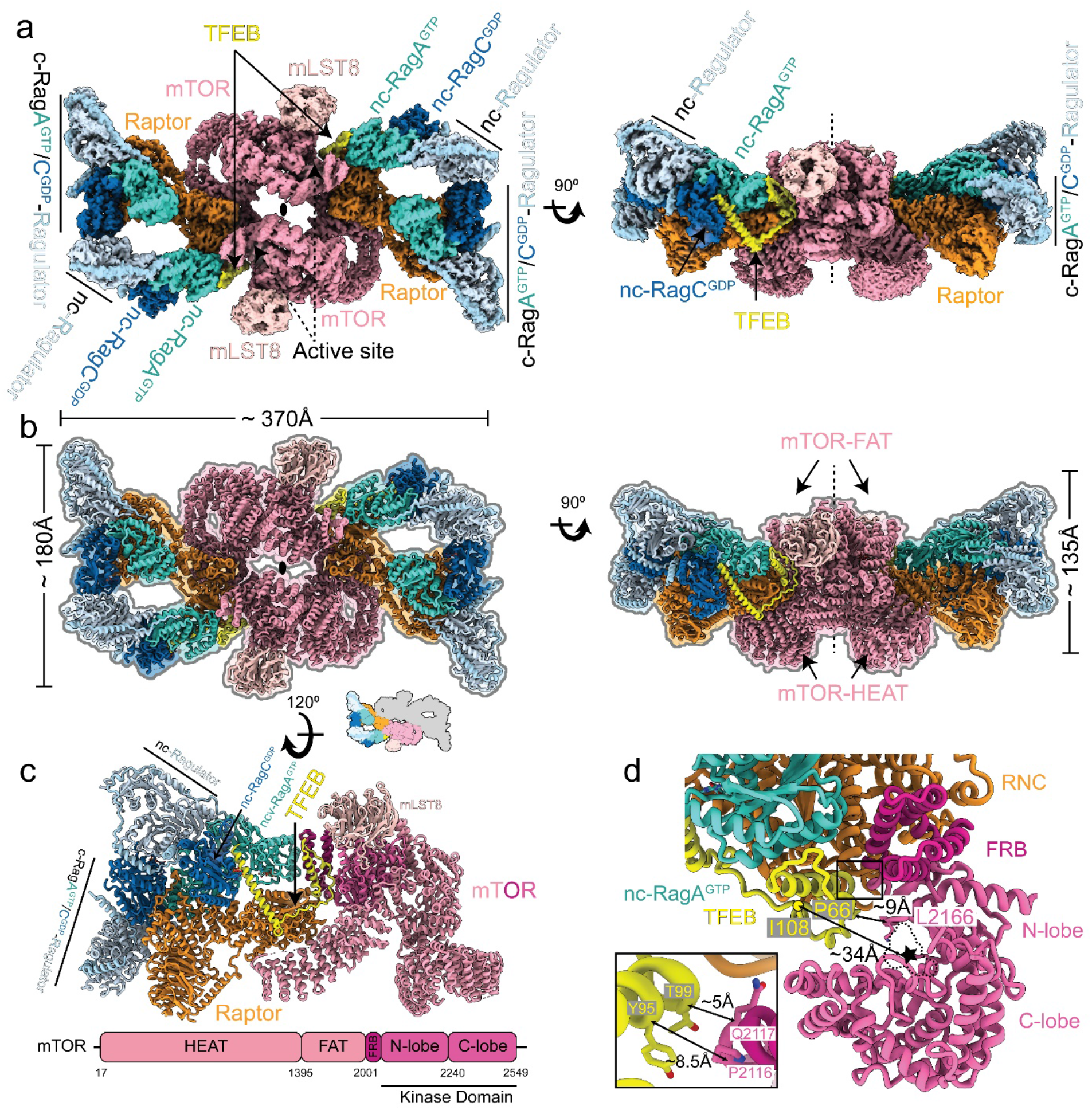
Architecture of the mTORC1-TFEB-Rag-Ragulator megacomplex. **a**, Composite cryo-EM density map of the dimeric mTORC1-TFEB-Rag-Ragulator megacomplex shown from top and side views. The active sites of mTOR are labeled with dash arrows. The twofold axis is labeled as an oval symbol in the top view and a dash line in the side view. **b**, Atomic model of the dimeric megacomplex shown in the same orientation as in (**a**). **c**, The ribbon model of an asymmetric unit. The domain organization of mTOR is shown. **d**, Focused view of the active site of mTOR, the HEAT and FAT domains are omitted for clarity. The ATP binding site is outlined with a dash line. The distance between Cα atoms of Pro^66^ of TFEB and Lys^2166^ of mTOR is drawn with a double-headed arrow. The distance between Ile108 of TFEB and the active site of mTOR is calculated based on the distance between Cα atoms of Ile^108^ of TFEB and Asp^2338^ of mTOR. The inset highlights the close contact between TFEB and the hinge loop (residues 2115-2118) at the end of mTOR FRB domain. Distances are calculated based on the Cα atoms.

The long axis of the megacomplex is about 370 Å, and displays a curved geometry, with the HEAT domain and FAT domain of mTOR facing the convex and concave side, respectively. The binding mode of TFEB to the Rags and Raptor is essentially unchanged in the presence of the entire mTORC1 complex. Residues 2-108 of TFEB were resolved in the cryo-EM structure, essentially as before, with the rest of TFEB not visualized despite the presence of a non-hydrolyzable ATP analogue and the presence of sequences containing phosphorylation sites. The inability to visualize these regions may be due to inherent flexibility in the sequences containing phosphorylation sites. The Pro-loop and α2 of TFEB are positioned near the active site of mTOR, surrounded by the FKBP12-rapamycin-binding (FRB) domain, the kinase domain (KD) N lobe and C lobe (Fig. 5c). A limited direct contact occurs between TFEB and mTOR, between Tyr^95^, Thr^99^ of TFEB and a hinge loop (residues 2115-2118) at the end of mTOR FRB domain (Fig. 5d)

The structure places the phosphorylation sites at Ser122 and Ser142 of TFEB close enough to reach the active site of mTOR. The last ordered residue of TFEB is ~34 Å from the catalytic residue Asp^2338^ of mTOR (Fig. 5d). This places even the most N-terminal of the sites, Ser122, close enough for delivery to the active site. In principle, the predicted flexible sequence from 109-210 could be long enough to deliver Ser211 to the second (distal) active site in the mTOR dimer, which is ~116 Å (linear distance) from TFEB residue 108. TFEB phosphorylation by mTORC1 is independent of Rheb ^25^. In the case of Rheb-dependent mTORC1 activation, a large-scale conformational change of mTORC1 was observed in the Rheb-bound state, which is thought to be essential for phosphorylation of TOS-containing substrates ^29^. Our structure shows how the Rheb-dependent conformational change in mTORC1 is unnecessary for TFEB phosphorylation, consistent with the Rheb-independence of TFEB as a substrate ^25^.

## Discussion

The structures described above provide a structural confirmation of the hypothesis that TFEB is regulated by its presentation to mTORC1 by a uniquely elaborate RagC^GDP^-dependent complex ^25^. The RagC^GDP^ state is required, in the first instance, to maintain favorable contacts with the TFEB N-terminal helix in the non-canonical Rag dimer. In the canonical Rag dimer, the RagC^GDP^ state is also important, in this case, for recruitment of the non-canonical Ragulator to the rest of the complex. The surprisingly elaborate complex involved in presenting TFEB for phosphorylation by mTORC1 thus depends on two molecules of RagC^GDP^. This mechanism serves to increase the stringency of TFEB regulation by the RagC GAP, FLCN. In the past few years, it has emerged that FLCN controls the phosphorylation of TFEB and other MiTF transcription factors, but not that of many other well-known mTORC1 substrates such as S6 kinase, ULK1, and 4E-BP1 ^25,26,33,34^. As the tumor suppressor whose mutation is responsible for Birt-Hogg-Dubé syndrome ^35^, FLCN is under exceptionally tight regulation. An elaborate set of structural gymnastics keep FLCN inactive by retaining it in the lysosomal folliculin complex (LFC) under starvation conditions ^26^, and reactivating it via inter-cleft competition with nutrient-activated SLC38A9 upon refeeding ^27^. To this picture of stringent regulation of FLCN, we have now added an even more elaborate and stringent mode of regulation of TFEB phosphorylation downstream of FLCN.

The structures described here confirmed several predictions in the literature, while also revealing many unexpected features. The N-terminal 30 amino acids of TFEB were correctly predicted to be essential for its Rag binding, lysosomal localization, and mTORC1 phosphorylation^30^. The role of the subsequent 80 residues, however, was unanticipated. The interaction of Raptor with the canonical Rag-Ragulator complex was as expected^22,23^. The presence of direct TFEB-Raptor interactions, and the existence of a second Rag-Ragulator complex in the presence of TFEB were also unexpected. The data presented here leave open the question as to whether the direct TFEB-Raptor interactions are important for function. The TFEB-Rag interactions are both extensive and essential, and the TFEB-Rag-Ragulator complex is biochemically stable in the absence of Raptor. Thus, it is still uncertain if the direct TFEB-Raptor interface is required for function, despite its contribution to structural ordering. Finally, the interaction between the G-domain of the canonical RagC with the non-canonical Ragulator is the first documentation of a direct structural interaction between a Rag G-domain and Ragulator.

Overall, this work provides structural evidence of a non-canonical mTORC1 signaling that allows selective control of TFEB activity under specific conditions ^25,39–41^. The hypophosphorylation and consequent hyperactivation of TFEB in the absence of FLCN drives increased mTORC1 activity and tumorigenesis in BHD syndrome ^25^. Thus, in BHD, there might be therapeutic benefits to enhancing TFEB phosphorylation by bypassing RagC^GDP^. The complexity and stringency of the structural mechanism for RagC^GDP^-dependent phosphorylation suggests this will be challenging. On the other hand, enhanced activation of TFEB may be desirable in treating lysosomal storage disorders ^7,8^, promoting clearance of toxic aggregates and debris in neurons ^9–12^, and preventing nonalcoholic fatty liver disease through lipid clearance ^42^.

The structure presented here identifies multiple novel interfaces that could be targeted to such an end.

## Acknowledgements

We thank S. Fromm and T. Stasyk for contributions to early stages of the project, and J. Remis and D. Toso for cryo-EM facility support. This work was supported by Genentech as part of the Alliance for Therapies in Neuroscience and the National Institute of General Medical Sciences, NIH, R01 GM111730 (J.H.H.), the Italian Telethon Foundation (to G.N. and A.B.), Associazione Italiana per la Ricerca sul Cancro A.I.R.C. (MFAG-23538 to G.N.; IG-22103 and 5×1000-21051 to A.B.), MIUR (PRIN 2017YF9FBS to G.N.; PRIN 2017E5L5P3 to A.B.) and the European Research Council H2020 AdG (LYSOSOMICS 694282 to A.B.).

## Data availability

Structural coordinates will be deposited in the RCSB and EM density maps in the EMDB prior to publication.

## Methods

### Protein expression and purification

The full-length codon-optimized human TFEB with S211A and R245-247A mutations, human RagC with S75N mutation, and human RagA with Q66L mutation were synthesized (Twist Bioscience) and cloned into a pCAG vector individually. The TFEB (S211A, R245-247A) construct included a TEV-cleavable GFP-His_10_ tag at the C-terminus. The RagC (S75N) construct included a TEV-cleavable GST tag at the N terminus, while the RagA (Q66L) was tagless. For the expression and purification of the TFEB-Rag GTPases complex, the HEK293F GnTI-cells were transfected with a total of 1mg plasmid DNA (333 μg TFEB,400 μg RagA, and 267 μg RagC) and 4 mg polyethylenimine (PEI) (Sigma-Aldrich) per liter at a density of 1.5–1.8 × 10^6^ cells/ml. Cells were collected after 48 hours, and lysed via gentle nutating in wash buffer (50 mM HEPES, 150 mM NaCl, 2.5 mM MgCl_2_,1mM TCEP, pH 7.4) supplemented with 0.4% CHAPS and Protease Inhibitor (Roche) for 1 hour. Lysate was cleared by centrifugation at 35,000 × g for 35 minutes. Supernatant was incubated with glutathione Sepharose 4B (GE Healthcare) resin for two hours. The resin was then first washed in the modified wash buffer with 200 mM NaCl and 0.3% CHPAS, and then in the wash Buffer. The complex was eluted from the resin by a wash buffer with 10 mM reduced glutathione, and then incubated with Tobacco Etch Virus (TEV) protease overnight. Eluted complexes were concentrated and further purified by size exclusion chromatography using a Superose 6 10/300 GL (GE Healthcare) column equilibrated in the wash buffer. All purification steps were performed at 4°C. Proteins were flash frozen in liquid nitrogen and stored in −80°C.

The human Ragulator complex (GST-TEV-Lamtor1, His6-TEV-Lamtor2) was expressed in *Spodoptera frugiperda* (Sf9) cells via baculovirus infection and purified as previously described ^37^. In brief, Sf9 cells were pelleted after 72 hours of baculovirus infection and lysed in the wash buffer with 1% Triton X-100 and protease inhibitor. The cleared supernatant after centrifugation was applied to Ni-nitrilotriacetic acid (NTA) gravity column (Thermo Scientific), washed with the wash buffer containing 200 mM NaCl, and eluted with the wash buffer with 250 mM imidazole. The elution was then applied to glutathione Sepharose 4B (GE Healthcare) gravity column, washed with the wash buffer. The complex was then eluted by on-column TEV-cleavage overnight without nutation. Further purification was done by SEC with a Superdex 200 10/300 GL column (GE Healthcare) column.

Three subunits of the human mTORC1 complex (mTOR, Raptor, mLST8) were codon-optimized and synthesized (GenScript). The mTOR gene was cloned into a pCAG vector without a tag, while Raptor and mLST8 were individually cloned into a pCAG vector with a uncleavable tandem 2×Strep II-1×FLAG-tag. The mTORC1 complex was produced in a similar manner as TFEB-Rag GTPases complex, except that the total amount of DNA was increased to 1.35 mg (900 μg mTOR, 250 μg Raptor, and 200 μg mLST8) per liter cells. The purification procedure of the mTORC1 complex is similar as previously described ^22^. Differently, Strep-Tactin resin (IBA Lifesciences) was used for the affinity purification and the complex was eluted with the wash buffer (50 mM HEPES, 150 mM NaCl, 1mM TCEP, pH 7.4) containing 10 mM D-desthiobiotin. The mTORC1 complex and free RAPTOR were separated by anion exchange chromatography on a 5 mL HiTrap Q column (GE Healthcare). The fractions containing mTORC1 complex and free Raptor were concentrated to 1.3 mg/mL and 0.5 mg/ml, respectively. Purified proteins were flash frozen in liquid nitrogen and stored at −80 °C.

### Cryo-EM sample preparation and imaging

The Raptor-TFEB-Rag-Ragulator complex was prepared by incubating 0.48 μM Raptor, 0.59 μM TFEB-Rag GTPases, 1.42 μM Ragulator, 9.5 μM GTP, and 9.5 μM GDP in the wash buffer on ice for 5 hours. Further purification of the complex was achieved by running a Superose 6 10/300 GL column. Fractions containing the fully assembled Raptor-TFEB-Rag-Ragulator complex were concentrated to 0.8 mg/ml for cryo-EM sample preparation.

The mTORC1-TFEB-Rag-Ragulator megacomplex was reconstituted in two steps. First, the TFEB-Rag-Ragulator complex was formed by incubating 2.5 μM TFEB-Rag GTPases, 7.4 μM Ragulator, 25 μM GTP, and 25 μM GDP in the wash buffer on ice for 1 hour. It is further purified through a Superose 6 10/300 GL column and concentrated to 1.2 mg/ml. And then, 0.36 μM mTORC1 complex, 1.8 μM TFEB-Rag-Ragulator complex, 18 μM GTP, 18 μM GDP, and 36 μM AMPPNP were incubated in 100 μL wash buffer containing 5 mM TCEP on ice for 5 hours. Assembled mTORC1-TFEB-Rag-Ragulator megacomplex was further concentrated to ~1 mg/ml for cryo-EM sample preparation.

Cryo-EM specimens were prepared by applying 3 μL of freshly reconstituted complex to a glow-discharged (PELCO easiGlow, 45 s in air at 15 mA and 0.4 mbar) holey carbon grid (C-flat: 2/1-3C-T) and vitrified using a FEI Vitrobot Mark IV (Thermo Fisher Scientific) after blotting for 3 seconds with blot force 18, 2 Whatman 595 papers on the sample side and 1 Whatman 595 paper on the back side at 6°C with 100% relative humidity.

Cryo-EM images of the Raptor-TFEB-Rag-Ragulator complex and mTORC1-TFEB-Rag-Ragulator megacomplex were recorded under a Titan Krios G3 microscope (Thermo Fisher Scientific) equipped with a Gatan Quantum energy filter (slit width 20 eV) and operated at 300 kV. Automated data acquisition was achieved using SerialEM ^43^ on a K3 Summit direct detection camera (Gatan) in the super-resolution CDS mode with a pixel size of 0.525 Å and a defocused range of −0.8 to −2.2 μm. Beam shift was enabled to encompass 4 exposures per hole and 9 holes per stage shift. The beam intensity was adjusted to a dose rate of ~1 e^-^ per Å^2^ per frame for a 50-frames movie stack with a total exposure time of 7.6 s. A total of 10,080 and 17,028 movie stacks were recorded for the Raptor-TFEB-Rag-Ragulator complex and the mTORC1-TFEB-Rag-Ragulator megacomplex, respectively.

Cryo-EM images of the TFEB-Rag-Ragulator complex were recorded under a Talos Arctica microscope (Thermo Fisher Scientific) operated at 200 kV. Automated data acquisition was achieved using SerialEM ^43^ on a K3 Summit direct detection camera (Gatan) in the superresolution CDS mode with a pixel size of 0.5575 Å and a defocused range of −0.8 to −2.2 μm. Beam shift was enabled to encompass 9 exposures per stage shift. The beam intensity was adjusted to a dose rate of ~1 e^-^ per Å^2^ per frame for a 50-frames movie stack with a total exposure time of 8.6 s. A total of 3,438 movie stacks were recorded for the TFEB-Rag-Ragulator complex.

### Cryo-EM data preprocessing

Super-resolution movie stacks were motion corrected and binned 2× by Fourier cropping using MotionCor2 with ^44^. Motion-corrected micrographs were primarily processed following the workflow in cryoSPARC v3 ^45^.

Data processing scheme for the Raptor-TFEB-Rag-Ragulator complex and mTORC1-TFEB-Rag-Ragulator megacomplex was shown in Extended data Fig. 2a and Extended data Fig. 8a, respectively. Due to the size of the datasets, micrographs were split and processed following the same protocol and then combined for homogeneous refinement. CTF determination was done by patch CTF in cryoSPARC v3. Blob picker and template picker were both used to maximize the number of initially picked particles. 2D classification was only used to remove obvious ‘junk’ particles (e.g., ice and chaperonin contaminants). Heterogeneous refinement following the ab initio reconstruction was used to select good particles, preserving potential rare views for those particles that could not be identified in 2D classification. After extensive cleaning using 2D classification and heterogeneous refinement, particles were merged, and the duplicates were removed with a 100-Å cutoff distance. Homogeneous refinement was then performed for the full dataset. Further cleaning of the full dataset was accomplished either by 3D classification with the “skip_align” option using RELION3 ^46^, or 3D classification function in cryoSPARC v3. The conversion of data files between cryoSPARC v3 and RELION3 was done using UCSF pyem ^47^. Local refinement was used to produce final cryo-EM maps for model building. For the mTORC1-TFEB-Rag-Ragulator megacomplex, symmetry expansion followed by local refinement was also used to generate the cryo-EM map of an asymmetric unit.

In summary, a 3.6-Å resolution map was obtained from 169,720 particles for the TFEB-Rag-Ragulator complex. Three local refinement maps were resolved for the Raptor-TFEB-Rag-Ragulator complex, including Raptor (377,569 particles), canonical Rag-Ragulator (377,569 particles), and non-canonical Rag-Ragulator (273,453 particles) to the resolutions of 2.8 Å, 2.9 Å, and 2.9 Å, respectively. For the mTORC1-TFEB-Rag-Ragulator megacomplex, two main populations containing either one (103,274 particles) or two copies (96,166 particles) of the TFEB and non-canonical Rag-Ragulator were both resolved with C1 symmetry to the resolution of 3.8 Å. Symmetry expansion and local refinement of the population with two copies of the TFEB and non-canonical Rag-Ragulator yielded a 3.2-Å resolution map.

The overall resolution of all these reconstructed maps was assessed using the gold-standard criterion of Fourier shell correlation ^48^ at 0.143 cutoff ^49^. Local resolution estimation ^50^ and local filtering were done within cryoSPARC v3.

### Atomic model building and refinement

To build the atomic model for Raptor-TFEB-Rag-Ragulator complex, we first fit the previous Raptor-Rag-Ragulator (PDB 6U62) structure in our cryo-EM map as rigid body using University of California San Francisco (UCSF) ChimeraX ^51^. The fragments of the TFEB model were initially obtained from AlfaFold2 prediction ^52^, and manually docked into our cryo-EM map. A composite map combining the three focused refinement maps was assembled using PHENIX ^53^. Model refinement against the composite map was performed by real-space refinement in PHENIX ^54^. Manual model building was done with COOT ^55^ and ISOLDE ^56^ to inspect and improve local fitting. The iterative process of refinement and the manual building was conducted to achieve the best model. For the mTORC1-TFEB-Rag-Ragulator megacomplex, the refined Raptor-TFEB-Rag-Ragulator and previous mTOR (PDB 6BCX) structures were docked in our cryo-EM map. A composite map of the symmetric complex using the focused refinement asymmetric unit was generated. The same model building procedure was performed as described above. All the figures and movies were made using UCSF ChimeraX.

### Materials and plasmids for cellular assays

Reagents used in this study were obtained from the following sources:

Antibodies to mTOR (Cat# 2983 - 1:100 IF) Phospho-p70 S6 Kinase (Thr389) (1A5) (Cat# 9206 - 1:1000 WB), p70 S6 Kinase (Cat# 9202 - 1:1000 WB), 4E-BP1 (Cat# 9644 - 1:1000 WB), Phospho-4E-BP1 (Ser65) (Cat# 9456 - 1:1000 WB), TFEB (Cat# 4240 - 1:1000 WB), Phospho-TFEB S211 (Cat# 37681 – 1:1000 WB) were from Cell Signaling Technology; antibodies to GAPDH (6C5) (Cat# sc-32233 - 1:15000 WB) and LAMP-1 (H4A3) (Cat# sc-20011 - 1:500 IF) were from Santa Cruz; antibody to HA.11 Epitope Tag (Cat# 901513) was from Biolegend; HRP-conjugated secondary antibodies to Mouse (Cat# 401215 - 1:5000 dilution) and Rabbit (Cat# 401315 - 1:5000 dilution) IgGs were form Calbiochem.

Chemicals: Torin 1 (Cat# 4247) was from Tocris; Protease Inhibitor Cocktail (Cat# P8340) and puromycin (Cat# P9620) were from Sigma Aldrich; PhosSTOP phosphatase inhibitor cocktail tablets (Cat# 04906837001) were from Roche.

Plasmids: The plasmid encoding full–length TFEB-GFP was previously described (Settembre et al., 2011). pRK5-HA GST RagC wt (#19304) was a kind gift from David Sabatini (Addgene plasmid). All the mutants used in these cellular assays were generated by using QuikChange II-E Site-Directed Mutagenesis Kit (#200555, Agilent Technologies).

### Cell cultures

HeLa cells were cultured in MEM (Cat# ECB2071L, Euroclone) supplemented with 10% inactivated FBS (Cat# ECS0180L, Euroclone), 2 mM glutamine (Cat# ECB3000D, Euroclone), penicillin (100 IU/mL) and streptomycin (100 μg/mL) (Cat# ECB3001D, Euroclone) and maintained at 37°C and 5% CO2. RagC-KO HeLa cells were previously generated and described (PMID: 32612235). Cell lines were validated by morphological analysis and routinely tested for absence of mycoplasma.

### Cell treatments

For experiments involving amino acid starvation, cells were rinsed twice with PBS and incubated for 60 min (unless stated otherwise) in amino acid-free RPMI (Cat# R9010-01, USBiological) supplemented with 10% dialyzed FBS. Serum was dialyzed against 1x PBS through 3500 MWCO dialysis tubing to ensure absence of contaminating amino acids. For amino acid re-feeding, cells were re-stimulated for 30 min with 1X water-solubilized mix of essential (Cat#11130036, Thermo Fisher Scientific) and non-essential (Cat# 11140035, Thermo Fisher Scientific) amino acids resuspended in amino acid-free RPMI supplemented with 10% dialyzed FBS, plus glutamine. Where reported, cells were incubated with 250 nM Torin1 during amino acid re-stimulation.

### Cell lysis and western blotting

Cells were rinsed once with PBS and lysed in ice-cold lysis buffer (250 mM NaCl, 1% Triton, 25mM HEPES pH 7.4) supplemented with protease and phosphatase inhibitors. Total lysates were passed 10 times through a 25-gauge needle with syringe, kept at 4°C for 10 min and then cleared by centrifugation in a microcentrifuge (14,000 rpm at 4°C for 10 min). Protein concentration was measured by Bradford assay. Cell lysates were resolved by SDS-polyacrylamide gel electrophoresis on 4%–12% Bis-Tris gradient gels (Cat# NP0323PK2 NuPage, Thermo Fischer Scientific) and analyzed by immunoblotting with the indicated primary antibodies.

### Confocal microscopy

Cells were grown on 8-well Lab-Tek II - Chamber Slides, treated as indicated, and fixed with 4% paraformaldehyde (PFA) for 10 minutes at RT. Blocking was performed with 3% bovine serum albumin in PBS + 0.02% saponin for 1 hour at RT. Immunostainings were performed upon dilution of primary antibodies in blocking solution and overnight incubation at 4C, followed by three washes and secondary antibody incubation in blocking solution for 1 hour at RT. After additional three washes, coverslips were finally mounted in VECTASHIELD^®^ mounting medium with DAPI and analyzed using LSM 800 or LSM 880 + Airyscan systems (Carl Zeiss), with a Plan-Apochromat 63×/1.4 NA M27 oil immersion objective using immersion oil (#518F, Carl Zeiss) at room temperature. The microscopes were operated on the ZEN 2013 software platform (Carl Zeiss). After calculation of processing for the airyscan, images were processed in the ImageJ 1.47v. Mander’s Co-localization Coefficients (MCC) was calculated using JACoP ImageJ Plugin.

### Immunoprecipitation

HeLa cells stably expressing TFEB-GFP were previously described ^13^. Cells grown on 10cm culture dishes were transiently transfected with the different HA-GST-RagC mutants and wildtype HA-GST-RagA using Fugene HD (#E2311, Promega). The following day, cells were treated with 330nM Torin I (# 4247, Tocris) for 1 hour. At the end of the incubation period, the cells were washed twice with ice cold PBS and lysed in 25mM HEPES pH7.4, 250mM NaCl, 1% Triton supplemented with protease and phosphatase inhibitors. Lysates were passed 5 times through a 25-gauge needle with syringe and then cleared by centrifugation (14,000 rpm at 4°C for 10 min). Lysates were then incubated with HA beads (#A2095, Sigma) at 4°C for 2 hours, washed with 40x the beads volume of lysis buffer, and eluted from the beads. Aliquots of the lysates and eluates were resolved by SDS-polyacrylamide gel electrophoresis on 8%, 10% or 15% SDS-PAGE gels and analyzed by immunoblotting with the indicated primary antibodies.

