## Extended Data for "Structural basis for mTORC1-dependent regulation of the lysosomal and autophagic transcription factor TFEB"

### Cryo-EM data collection, refinement and validation statistics

| | Raptor-TFEB-Rag-Ragulator (EMDB-xxxx) (PDB xxxx) | mTORC1-TFEB-Rag-Ragulator (EMDB-xxxx) (PDB xxxx) |
|---|---|---|
| **Data collection and processing** | | |
| Magnification | 81,000 | 81,000 |
| Voltage (kV) | 300 | 300 |
| Electron exposure (e–/Å$^2$) | ~50 | ~50 |
| Defocus range (μm) | -0.8 to -2.2 | -0.9 to -2.1 |
| Pixel size (Å) | 1.05 | 1.05 |
| Symmetry imposed | C1 | C2 |
| Initial particle images (no.) | 3,768,278 | 5,505,615 |
| Final particle images (no.) | 273,453 | 96,166 |
| Map resolution (Å) | 3.1 overall (2.8/2.9/2.9 local refine) | 3.2 |
| FSC threshold | 0.143 | 0.143 |
| Map resolution range (Å) | ~2.6 to ~13.8 | ~2.6 to ~32 |
| **Refinement** | | |
| Initial model used (PDB code) | 6U62 | 6BCX |
| Model resolution (Å) | 3.0 | 3.4 |
| FSC threshold | 0.5 | 0.5 |
| Map sharpening *B* factor (Å$^2$) | -110.5 | -86.3 |
| Model composition | | |
| Non-hydrogen atoms | 27,280 | 95,250 |
| Protein residues | 3,423 | 11,922 |
| Ligands | 6 | 14 |
| *B* factors (Å$^2$) | | |
| Protein | 34.32 | 105.63 |
| Ligand | 18.95 | 73.09 |
| R.m.s. deviations | | |
| Bond lengths (Å) | 0.004 | 0.005 |
| Bond angles (°) | 0.724 | 0.877 |
| Validation | | |
| MolProbity score | 1.54 | 1.28 |
| Clashscore | 3.91 | 1.78 |
| Poor rotamers (%) | 0.00 | 0.03 |
| Ramachandran plot | | |
| Favored (%) | 94.68 | 95.01 |
| Allowed (%) | 5.32 | 4.86 |
| Disallowed (%) | 0.00 | 0.14 |

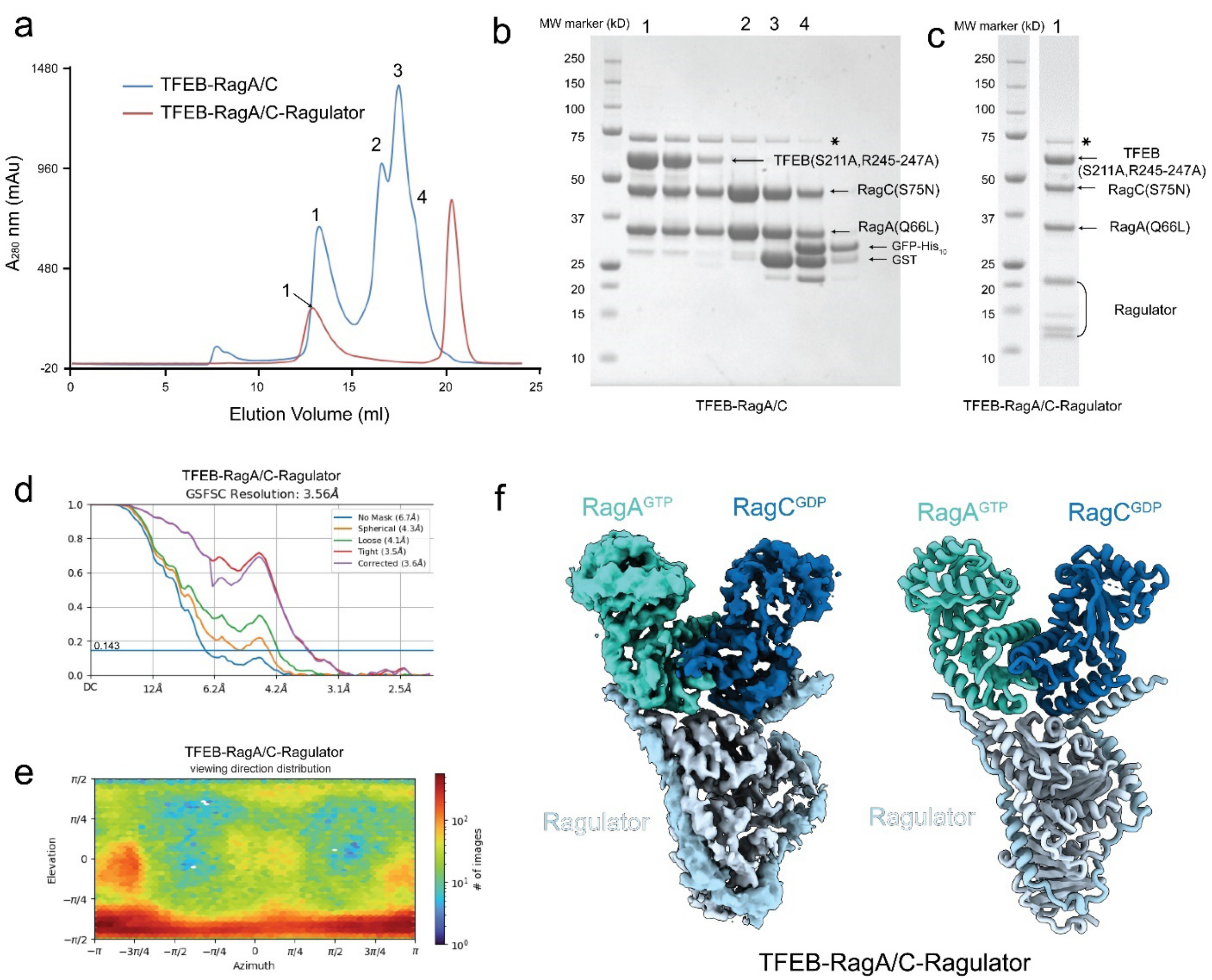

**Extended Data Fig.1 Purification, reconstitution, and cryo-EM structure determination of the TFEB-Rag-Ragulator complex.**

**a,** Gel filtration chromatography of the TFEB-Rag and TFEB-Rag-Ragulator complexes, using a Superose 6 10/300 GL (GE Healthcare) column. Corresponding peaks are labelled and analyzed by Coomassie blue staining SDS-PAGE in **b** and **c** for TFEB-Rag and TFEB-Rag-Ragulator complexes, respectively. **d-f,** cryo-EM structure determination of TFEB-Rag-Ragulator complex. **d,** Resolution estimation based on gold standard FSC. **e,** Orientation distribution of the reconstructed cryo-EM map. **f,** Side-by-side view of the cryo-EM density map and atomic model of the TFEB-Rag-Ragulator complex. Cryo-EM density for TFEB is not resolved. Asterisks in **b** and **c** indicate HSP70 contamination.

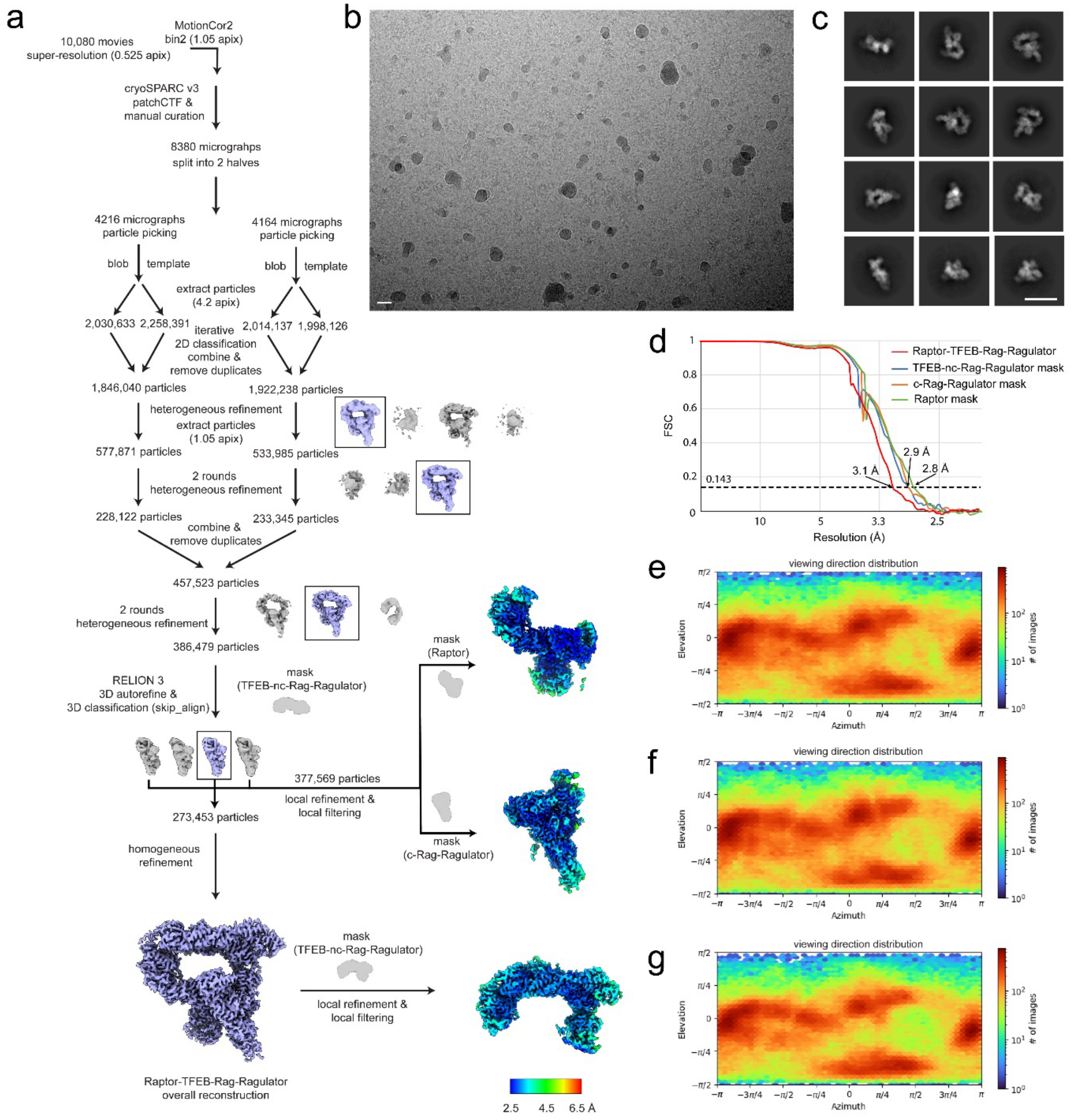

**Extended Data Fig. 2 Cryo-EM workflow of the Raptor-TFEB-Rag-Ragulator complex.**

**a,** Cryo-EM data processing diagram of the Raptor-TFEB-Rag-Ragulator complex. **b,** A representative micrograph of the dataset after motion correction. **c,** Selected 2D class average images showing different orientations of the complex. **d,** Resolution plots of the cryo-EM reconstructions with different masks. Orientation distribution of reconstructed cryo-EM maps with Raptor, c-Rag-Ragulator, and nc-Rag-Ragulator masks are shown in **e, f,** and **g,** respectively. Scale bars in **b** and **c** represent 20 nm.

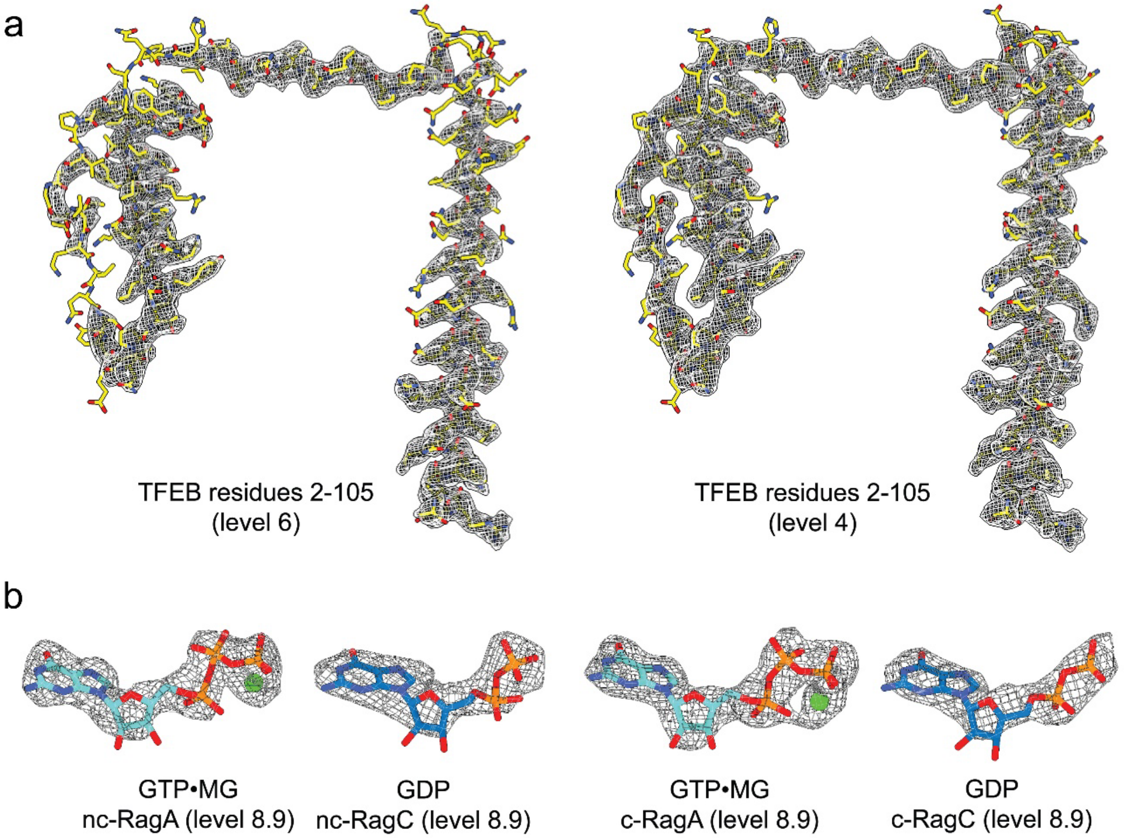

**Extended Data Fig. 3 Representative cryo-EM density of the Raptor-TFEB-Rag-Ragulator complex.**

**a,** Cryo-EM density of TFEB (2-105) at contour level 6 (left) and level 4 (right). **b,** Cryo-EM density of the nucleotides at contour level 8.9

a

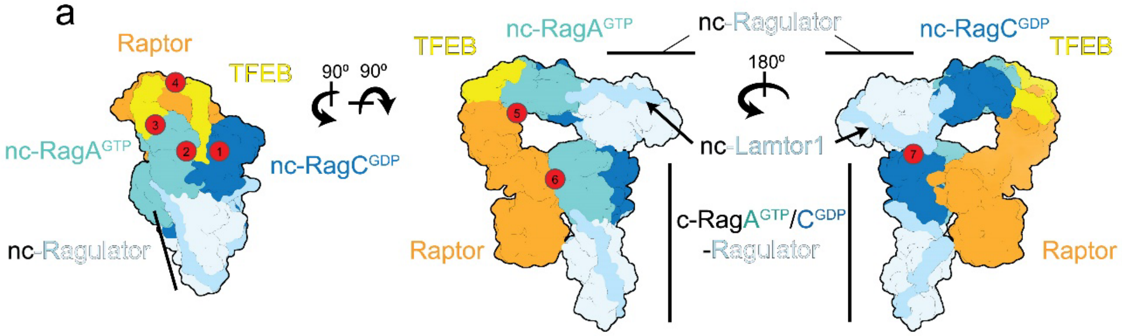

b

| Subunit 1 | Subunit 2 | Interaction interface Å² | |
|---|---|---|---|
| TFEB | nc-RagC$^{GDP}$ | ~570 | 1 |
| | nc-RagA$^{GTP}$ (inter-Rag G domain cleft) | ~580 | 2 |
| | nc-RagA$^{GTP}$ (outer-Rag G domain) | ~700 | 3 |
| | Raptor | ~1280 | 4 |
| Raptor | nc-RagA$^{GTP}$ | ~300 | 5 |
| | c-RagA$^{GTP}$ | ~1500 | 6 |
| nc-Lamtor1 | c-RagC$^{GDP}$ | ~380 | 7 |

**Extended Data Fig. 4 Interaction surface aera estimation of the Raptor-TFEB-Rag-Ragulator complex.**

**a,** A cartoon representation of the complex, labelled with red circles indicating different interaction surface. **b,** A table showing the interaction surface aera labelled in **a,** estimated by PISA.

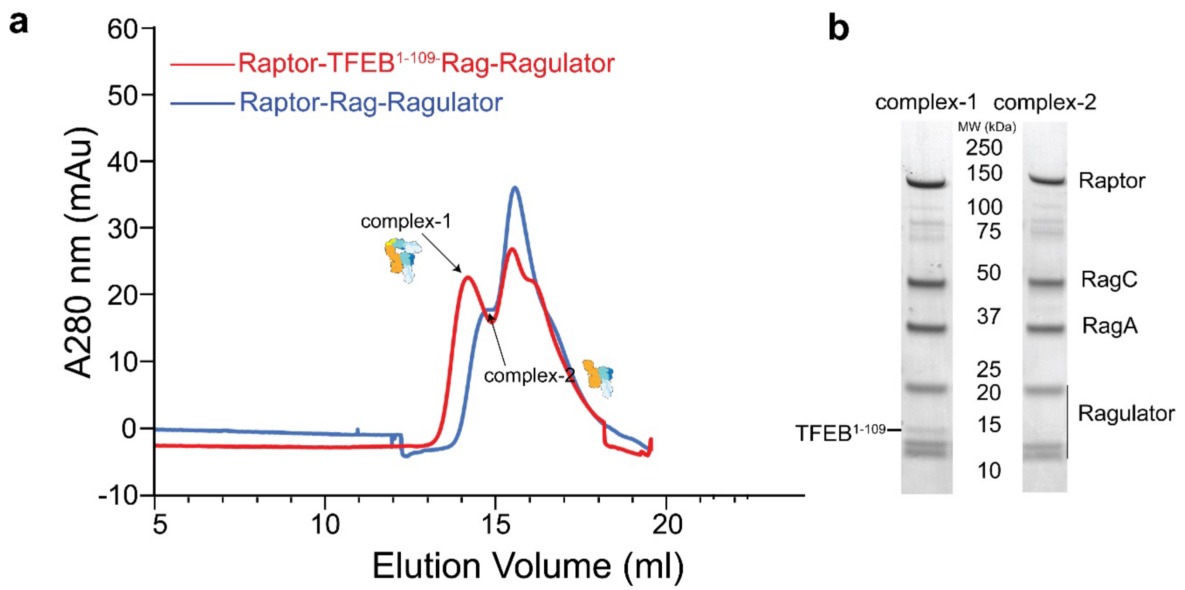

**Extended Data Fig. 5 Assembly of Raptor-TFEB$^{1-109}$-Rag-Ragulator complex**

**a,** Gel filtration chromatography of the Raptor-Rag-Ragulator and Raptor-TFEB$^{1-109}$-Rag-Ragulator complexes, using a Superose 6 10/300 GL (GE Healthcare) column. **b,** The peaks corresponding to the largest complex are analyzed by SDS-PAGE and stained by Coomassie blue.

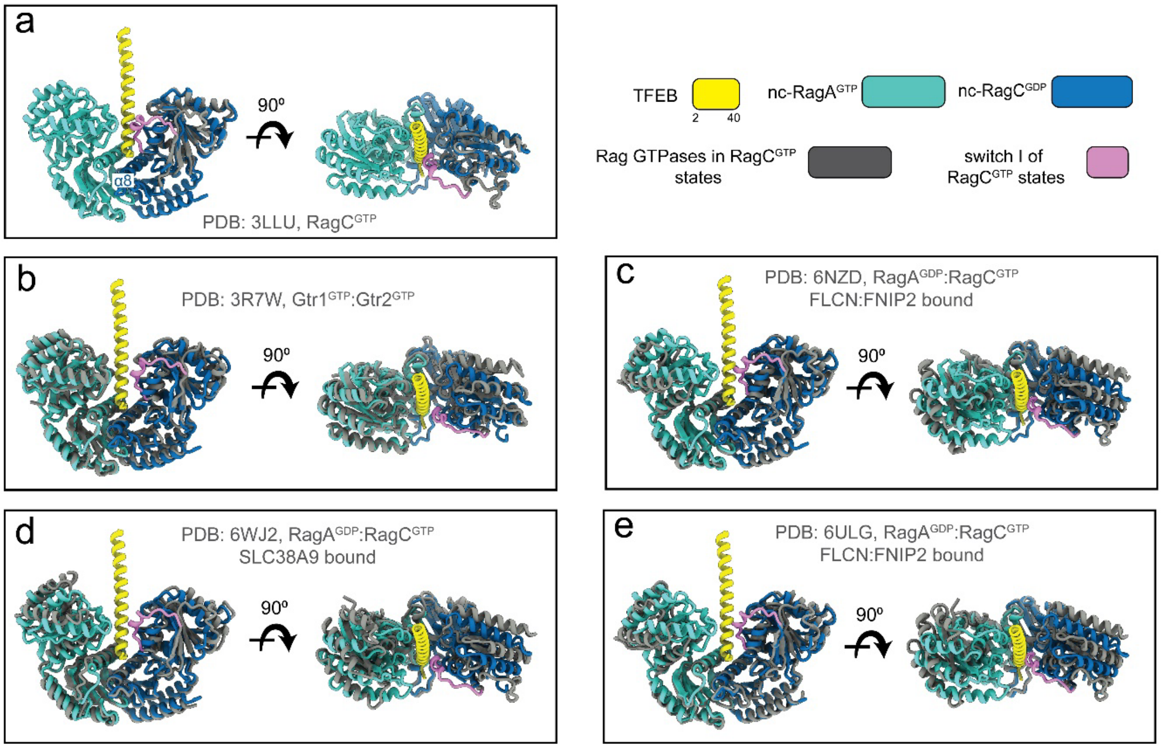

**Extended Data Fig. 6 Structural comparison between the active nc-Rag GTPases ($RagA^{GTP}$:$RagC^{GDP}$) and RagC in GTP-bound or empty states.**

Structures are superimposed based on the α8 of RagC. The nc-Rag GTPases are colored as in Fig.1. Structures of RagC in GTP-bound states are colored as gray, while the switch I regions are colored as pink. Residues 41-105 of TFEB are omitted.

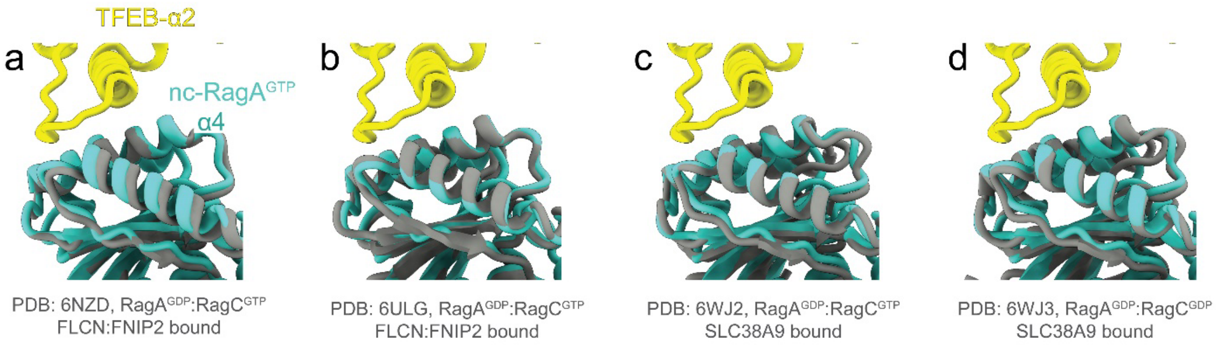

**Extended Data Fig. 7 Structural comparison between the nc-Rag$^{GTP}$ and RagA in GDP-bound states at the second TFEB contact site**

Structures are superimposed based on the α4 of RagA. The nc-RagA$^{GTP}$ and TFEB are colored as in Fig.1. Structures of RagA at GDP-bound states are colored as gray.

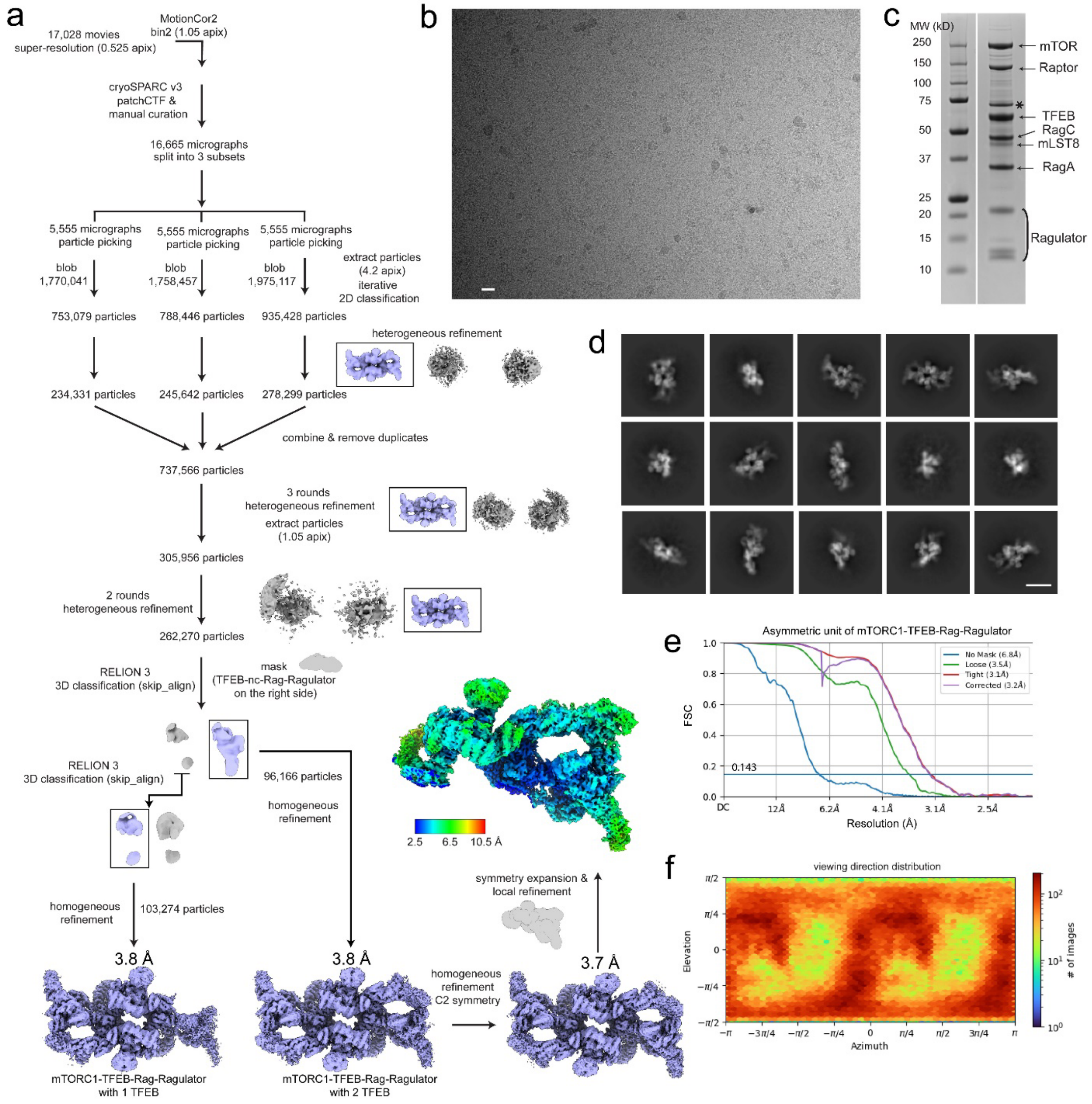

**Extended Data Fig. 8 Cryo-EM workflow of the mTORC1-TFEB-Rag-Ragulator megacomplex**

**a,** Cryo-EM data processing diagram of the mTORC1-TFEB-Rag-Ragulator megacomplex. **b,** A representative micrograph of the dataset after motion correction. **c,** SDS-PAGE of the reconstituted megacomplex stained by Coomassie blue. Asterisk indicates HSP70 contamination. **d,** Selected 2D class average images showing different orientations of the complex. **e,** Resolution plots of the asymmetric unit of mTORC1-TFEB-Rag-Ragulator megacomplex. **f,** Orientation distribution of reconstructed mTORC1-TFEB-Rag-Ragulator cryo-EM map. Scale bars in **b** and **d** represent 20 nm.

**Movie S1**: Atomic model of mTORC1-TFEB-Rag-Ragulator megacomplex.

Overview of the model built upon the cryo-EM density, highlighting novel interaction interfaces discussed in the manuscript.
